# Formation of DNA duplexes in the presence of urea as a chaotropic agent

**DOI:** 10.1101/2025.11.07.686900

**Authors:** Marcello Gottschalk, Robin Jacobi, Sophia Rosencrantz, Ruben R. Rosencrantz, Oleh Fedorych

## Abstract

Hybridization of fluorescent molecular probes, such as molecular beacons or linear molecular probes, with their molecular targets can be confirmed through fluorescence spectra. In this study, we investigate the interaction between fluorescent probes, specifically linear molecular probes, and their targets in the presence of chaotropic agents. Our findings show that double-stranded structures containing mismatched nucleotides (e.g., single-nucleotide polymorphisms) occupy lower energy levels (red shifted) compared to those without mismatched nucleotides. Molecular duplexes formed in buffers without chaotropic agents do not exhibit significant differences, independently on whether they contain mismatched nucleotides or are formed with perfectly matching targets. This effect appears in the presence of a naturally occurring chaotropic agent, urea, and was confirmed in the concentration range of 1 mM to 4 mM. These findings suggest that urea and similar agents may play a significant role in the formation of mismatched nucleotide structures.

## Introduction

In our previous work [1], we reported a method to detect changes in the fluorescence spectra of molecular beacons in their single and double stranded states. These changes were attributed to a reduction in the free energy of the molecular beacon upon hybridization with a complementary strand. Although molecular beacons are excellent candidates for diagnostic applications, they are very demanding in handling [2]. For instance, to make a molecular beacon accessible for hybridization, its stem must be destabilized and its loop unfolded which is mostly performed thermally or by chemical means. These conditions are difficult, if not impossible, to achieve in living systems. From this perspective, linear molecular probes (LMP), which lack secondary structure, offer a more practical alternative. LMP [3] are simple, single-stranded nucleic acid sequences that retain their high specificity for complementary targets. Unlike molecular beacons, LMP remain accessible for hybridization under physiological conditions, making them especially useful when working with unaltered or unprocessed biological samples. Notably, LMP are the key component of fluorescence in situ hybridization (FISH) [4], a widely used technique in cellular and molecular biology. Like molecular beacons, LMP used are fluorescently labeled and specifically bind to complementary DNA or RNA sequences, allowing visualization of genetic material under a fluorescence microscope. Despite decades of refinement, FISH as a method, still struggles with detecting single nucleotide polymorphisms (SNP) [5]. While the method yields quantitative data, it does not provide insight into the energetic dynamics of hybridization or the influence of the local environment on probe – target interactions. A method that combines both energetic and structural information would be particularly interesting [6].

We previously reported [1] that, in a highly chaotropic buffer, a molecular beacon hybridized with a mismatched target containing a single nucleotide polymorphism can form a duplex with lower free energy than one formed with a perfectly matched target. Although those conditions differ substantially from physiological environments, our goal was to reproduce this effect under biologically relevant conditions, specifically using a naturally occurring chaotropic agent. One of the most interesting candidates is urea, which at molar concentrations is known to denature DNA and RNA, lowering their melting temperatures by 2.25□°C per molar and slowing renaturation [7, 8]. However, the effect of urea at biologically relevant concentrations [9] on the hybridization of DNA strands remains largely unexplored. Therefore, we examined its influence at concentrations ranging from 1□mM to 4□mM.

## Materials and Methods

Oligonucleotides were delivered by biomers.net (Ulm, Germany). The LMP used in the experiments, comprises of a 23-nt probe 5′-AGA CCA GAA GAT CAG GAA CTC TA-3’, labeled at the 5′-end with rhodamine 6G. Two target strands were prepared, one of which was a perfect match (PM) for the probe 5′-TAG AGT TCC TGA TCT TCT GGT CT-3′ whereas the other contained a mismatching (MM), central nucleotide (bold underlined), 5′-TAG AGT TCC TGA T**T**T TCT GGT CT-3′ and scramble strand (SS) as a negative control 5′-GCT ATG CGT ATT GTT CAC TTG TC-3′. The LMPs were diluted to a final concentration of 0.1□µM in 50□mM Tris-HCl buffer (pH 7.4), and one of the three target strands was added such that the LMP concentration remained unchanged, while the target concentration was 0.2□µM. The 1:2 ratio of LMP to target was chosen to promote complete hybridization of the LMPs, and the fluorescence was expected to arise primarily from the duplex state. Urea was used as the chaotropic agent, with concentrations ranging from 0□mM to 4□mM in 1□mM increments. After preparation, samples were heated to 37□°C and then allowed to cool to room temperature. All measurements were performed at room temperature, and all solutions were protected from ambient light throughout the experiment.

Fluorescence measurements were performed using a custom-built micro-fluorescence setup (Fig.□1a). Linear molecular probes (LMPs) diluted in the sample were excited using a 532□nm diode-pumped solid-state laser with an emission power of 5□mW. The laser beam passed through a 90:10 (transmission:reflection) beam splitter, yielding an effective excitation output of approximately 0.5□mW. The reflected beam was coupled into an optical fiber via a fiber optic coupler. The other end of the fiber, which was cut and polished, was immersed directly into the analyte solution.

**Fig 1.**
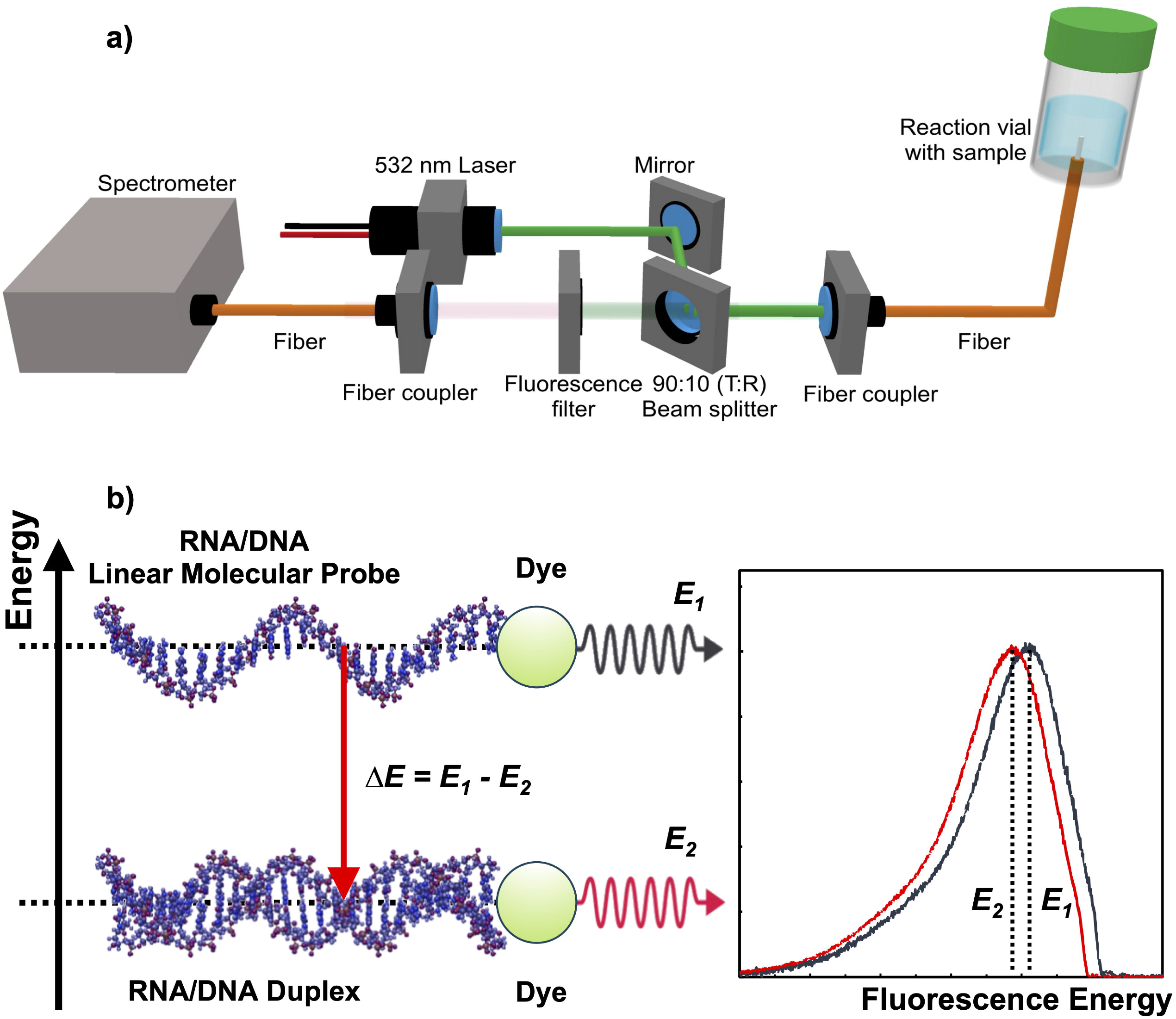
Experimental setup and method. **a)** Shema of custom-built micro-fluorescence setup used to measure the fluorescence of a linear molecular probe (LMP), with a bare fiber end immersed directly into the sample. **b)** Schematic representation of the principle: the LMP with an attached dye emits fluorescence with a peak energy E_1_ in its single stranded state. Upon hybridization with a target, the resulting duplex emits fluorescence with a peak energy E_2_. The energy difference ΔE = E_1_ – E_2_ corresponds to the hybridization energy associated with duplex formation.

The excitation light stimulated fluorescence emission from rhodamine 6G dye molecules covalently attached to the LMP. A portion of the emitted fluorescence was recaptured by the same optical fiber used for excitation. After passing through the beam splitter and a high-pass fluorescence filter, approximately 90% of the fluorescence signal was directed into a spectrometer via a second optical coupler. Typical measurements were performed for one minute, with each spectrum accumulated for one second displayed and stored on the control computer.

The setup operates in a near field configuration, allowing any changes in the refractive index of the sample, or between samples, to be considered negligibly small. The excitation power delivered to the samples was carefully adjusted to minimize potential heating effects caused by laser irradiation. For convenience in analysis, the fluorescence spectra were converted to energy units (eV), and a schematic representation of the measurement results is shown in Fig.□1b. In this schema, the LMP with an attached dye in its single stranded state occupies an energy level denoted as E_1_, and the dye emits a corresponding fluorescence spectrum. Upon hybridization with target nucleotides, the LMP forms a duplex that occupies a lower energy level (E_2_), where E_2_□<□E_1_. As a result, the fluorescence spectrum emitted by the rhodamine 6G dye is red-shifted relative to the single stranded state. The energy difference ΔE=E_2_ – E_1_ corresponds to the hybridization energy related with the formation of the duplex.

## Results and Discussion

At first the wavelength scale of the spectra was converted to eV for convenience [10]. Fluorescence intensity and its variations during the experiment were monitored solely as the indicator of sample quality. Even within the same batch of samples prepared from identical stock solutions, fluorescence intensity could vary by up to ∼50%. Therefore, the analysis focused primarily on the peak positions of the fluorescence spectra.

Direct observation of spectral shift, i.e., the decrease in fluorescence energy upon duplex formation compared to the single stranded state of the LMP, is not a trivial challenge. This challenge appears due to the small magnitude of the shift and the broad emission profile of the rhodamine 6G dye (see Fig.□2a). In this figure, normalized fluorescence spectra are shown for three conditions: LMP with a scrambled target (single stranded, blue line), LMP hybridized with a perfectly matching target (duplex, black line), and LMP hybridized with a target containing a single nucleotide polymorphism (duplex with SNP; red line). All measurements were performed in the presence of 2□mM urea.

**Fig 2.**
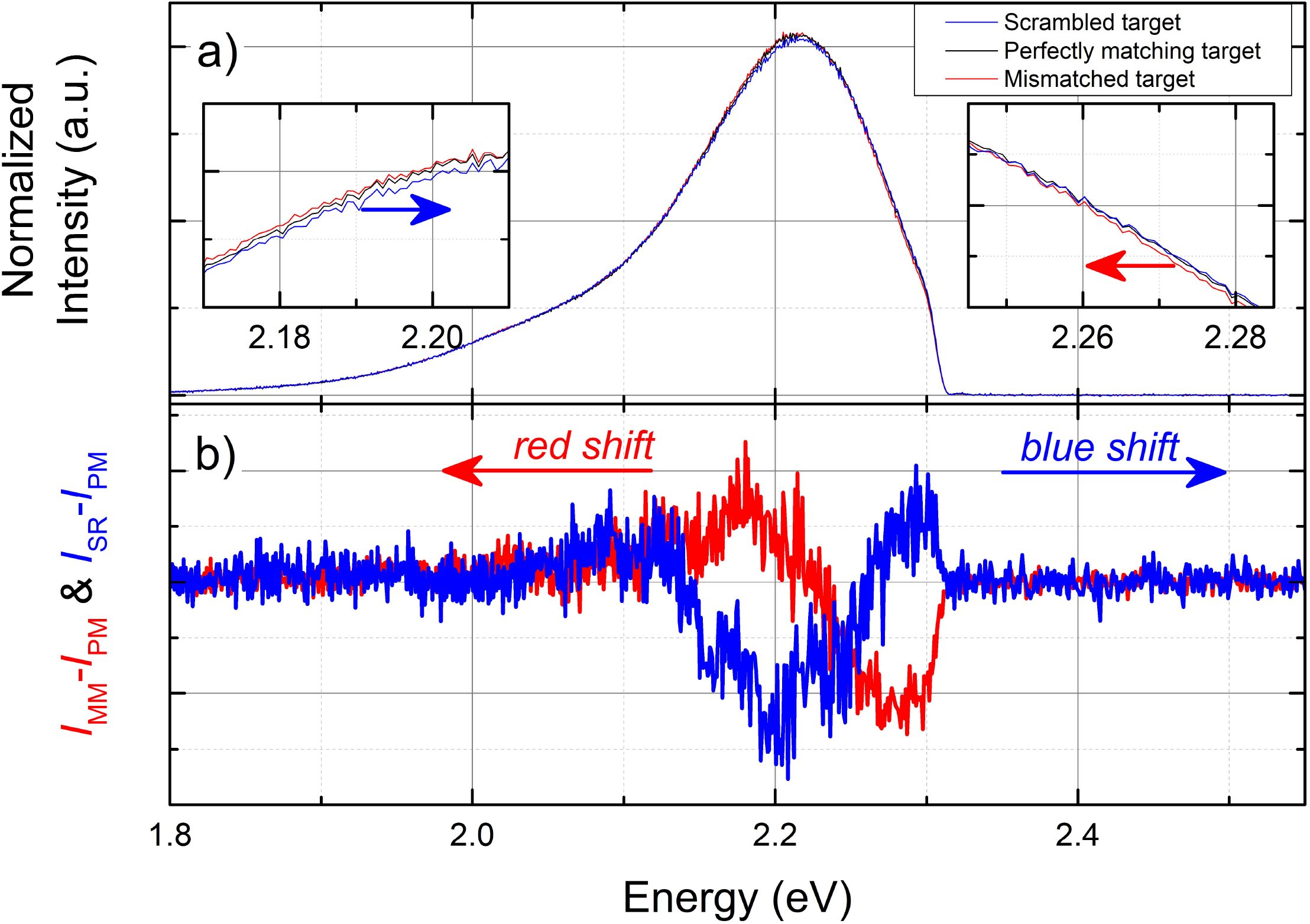
Fluorescence spectra and their energy shifts. **a)** Normalized fluorescence intensity spectra of rhodamine 6G dye attached to linear molecular probes (LMPs) measured in the presence of 2□mM urea under three conditions: with a scrambled target (blue line), hybridized with a perfectly matching target (black line), and hybridized with a target containing a single mismatched nucleotide (red line). The two zoomed insets display the energy displacement of the LMP spectrum measured with the scrambled target and the mismatched target in 2 mM urea-containing buffer. **b)** Two differential spectra obtained by subtracting the normalized spectrum of the perfectly matched duplex from the spectra of the scrambled (blue line) and mismatched (red line) targets. The color of each line corresponds to the direction of the spectral shift: blue indicates a shift to higher energy (blue shift), and red indicates a shift to lower energy (red shift).

To more clearly resolve the spectral shifts, differential spectra were computed by subtracting the normalized fluorescence spectrum of the perfectly matched duplex from the spectra of the other two conditions (Fig.□2b). These differential spectra show the direction of spectral shifts, either toward higher energy (blue shift) or lower energy (red shift). As seen, the differential spectrum for the scrambled target is blue shifted, while the duplex containing a polymorphic nucleotide exhibits a red shift relative to the perfect duplex.

We also performed a detailed analysis of all experimentally obtained spectra by fitting each with a sum of two Gaussian peak functions [11, 12]. One Gaussian peak corresponded to the emission of rhodamine 6G in its monomeric state, while the other represented emission from dye dimers. Although only the monomer emission is of practical interest, accurately determining the spectral peak position requires a model that accounts for both components. Therefore, all spectra from all samples were fitted using this double peak model. The resulting monomer peak positions were grouped according to three conditions: LMP with a scrambled target (single stranded state; blue points, Fig.□3a), LMP hybridized with a perfectly matching target (duplex; black points, Fig.□3b), and LMP hybridized with a target containing a single nucleotide polymorphism (SNP; gray and red points, Fig.□3c).

**Fig 3.**
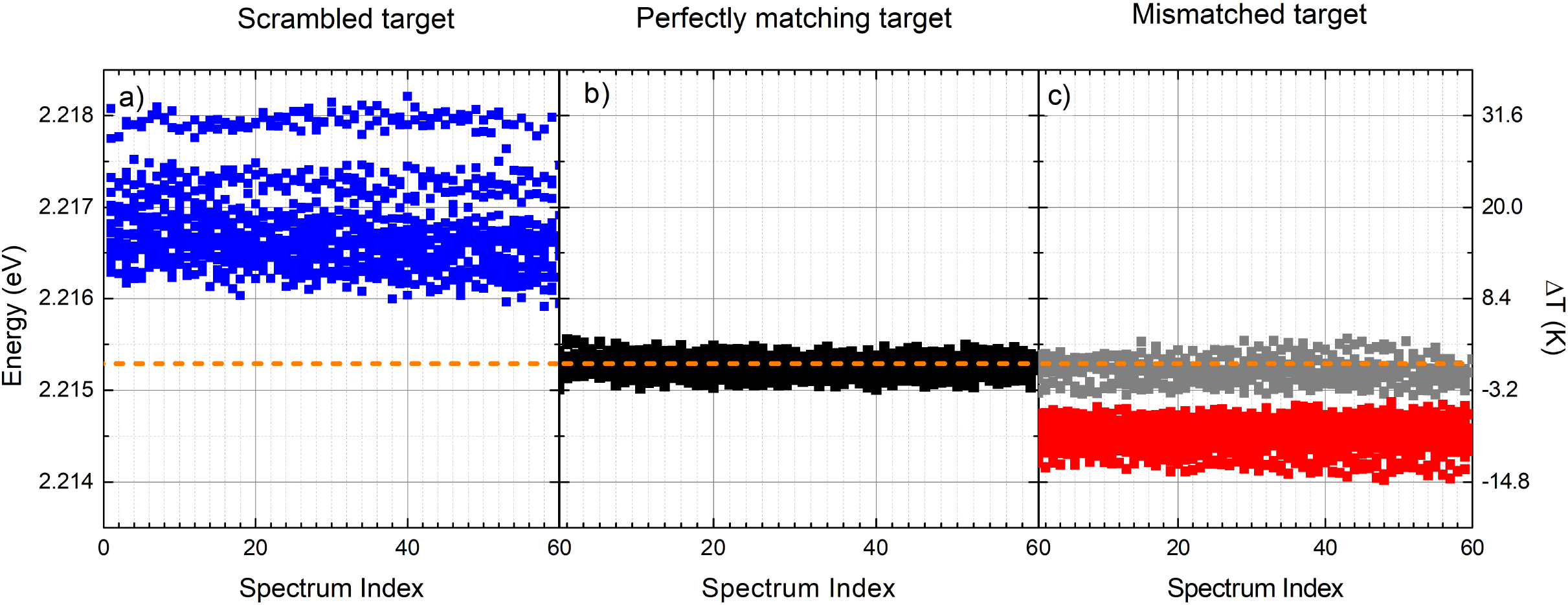
Emission energies of the LMP in the presence of three targets and varying urea concentrations. Fitted emission energies of the linear molecular probe from 60 spectra were measured under three conditions: **a)** single stranded state with a scrambled target (blue points), **b)** double-stranded state with a perfectly matching target (black points), and **c)** double stranded state with a target containing a single mismatched nucleotide. For the SNP duplex, the emission energies are distributed across two well resolved bands: one corresponding to samples without urea (gray points) and the other samples containing x mM urea (red points). The right axis displays a relative temperature scale, representing the thermal equivalent (in kelvins, K) of the energy change with respect to the reference energy level of the perfectly matched duplex.

The advantage of applying the fitting approach, particularly for broad and intense fluorescent emissions such as those exhibited by rhodamine 6G, is that it enables determination of the peak position with an accuracy higher the nominal resolution of the spectrometer. As shown in Fig. 2a, the rhodamine 6G monomer emission line, centered near 2.21□eV, exhibits a spectral width of approximately 0.1□eV. Therefore, for rhodamine 6G spectra measured with common spectrometers, peak fitting allows the line position to be determined with an accuracy around 0.1□meV.

The energy reported by the LMP in the case where it reacts with the scrambled target varies over more than 2□meV in total, or over 20□K in thermal equivalent (see Fig.□3a). We relate this large energy spread to internal nonuniformity within the samples, caused by the presence of two nucleic strands that are unable to hybridize and form a duplex structure, and therefore cannot reduce their total energy. As shown below, we do not observe any remarkable influence of urea on the energy in this case, which is not surprising for the strongly perturbed and unstructured solutions.

For the double stranded duplex formed with the perfectly matching target, the energy reported by the LMP falls within a band approximately 0.5□meV wide, corresponding to about 6□*K* in thermal energy (see Fig.□3b). Compared to the condition with the scrambled target, the band for the perfect duplex is narrower, but still relatively broad. This spread can be attributed to fluctuations in the concentrations of the LMP and the perfectly matching target between different samples. Similar to the previous condition, the presence of urea, despite its chaotropic nature, does not affect the energy of the LMP in the duplex state (see below). This is not surprising, as the duplex molecule represents the most stable and neutral state of DNA, with all hydrogen bonds fully engaged in stable pairing with the complementary strand. As a result, there are no unpaired bonds whose disruption could increase the duplex entropy in the presence of a chaotropic agent [13].

The final condition, where the LMP forms a double stranded structure with a single polymorphic nucleotide, demonstrates the most intriguing result. As shown in Fig.□3c, the emission energy of the LMP splits into two distinct bands, one corresponding to samples without urea, and the other to samples with urea. The energy levels for the mismatched samples without urea (gray points in Fig.□3c) overlap with those of the perfectly matched duplex and shows the similar band broadening. As can be seen from the figure, the values of the energy levels obtained for the duplexes with and without the SNP are very close. According to theoretical predictions [14], the energy of the mismatched duplex should be about 0.2 meV (2 K) higher than that of the perfectly matched duplex. Considering that the accuracy of peak position determination is around 0.1 meV, this is not sufficient to resolve the energy levels. Moreover, broadening of the energy levels, with a width of ∼ 0.5 meV (6 K), makes it even more difficult to resolve these levels.

The duplex containing a polymorphic nucleotide, in the presence of the chaotropic agent urea, exhibits a lower energy compared to both the same duplex without the chaotropic agent and the perfectly matched duplex [15-18]. The energy level broadening is approximately 0.7□meV (∼8□K), and the center of the band is shifted by about 0.*9*ΩmeV (∼11□K) relative to that of the perfectly matched duplex. Similarly, as above, we attribute the energy level broadening to variations in the concentrations of the LMP and target strands, particularly when at least half of the target molecules remain unpaired (i.e., unhybridized). However, the distinct and significant red shift observed for the less favorable duplex, relatively to the perfectly matched duplex, cannot be explained by classical nearest-neighbor theory [13], which predicts a blue shift of about 0.2 meV (2 K).

Our previous work reported a similar effect [1] using molecular beacons, guanidinium thiocyanate at higher concentrations, and distinct experimental conditions. This study employed low concentrations of chaotropic agent urea (max. 4 mM) and LMP, precluding significant duplex destabilization. Consequently, urea showed no observable impact on the perfectly matched duplex’s energy state, which exists in its most stable configuration, rendering it largely impervious to environmental effects.

The presence of a single mismatched nucleotide introduces an unpaired hydrogen bond, enabling interaction with the surrounding environment. This site allows the duplex to interact with water molecules, inducing their local ordering and a consequent reduction in entropy, thereby increase the free energy of the duplex. However, the introduction of a chaotropic agent, such as urea, disrupts the ordered water structure, leading to an increase in system entropy and a reduction in free energy. This so-called “hydrophobic contribution to the dissociation energy” [19] can yield to unexpected outcome. Under certain conditions, presence of the urea, thermodynamically less stable (mismatched) duplexes can become more energetically favorable than their perfectly matched duplexes, potentially resulting in mismatched duplexes preferential formation. Higher temperatures, however, reduce the dominance of this entropy driven hydrophobic effect, causing the system to revert to the expected behavior where more stable states (perfectly matched duplexes) exhibit higher melting temperatures than less stable mismatched duplexes.

Fig. 4 a) shows the experimentally measured dependence of the emission energy of the LMP on the concentration of urea under three conditions: single stranded state with the scrambled target (blue points, blue zone), double stranded state with the perfectly matching target (green points, gray zone), and double stranded state with the mismatched target (red points, red zone). As evident from the figure, the presence of urea has no remarkable influence on the energy of the LMP in either the single stranded state or the perfectly matched duplex. In contrast, the effect of urea on the mismatched duplex exhibits a binary character, the presence of urea activates the hydrophobic contribution, and after no further change in free energy is observed with increase of urea concentration. At lower concentrations of urea, one might expect a mixed state, where a portion of the duplexes containing SNPs are affected by the urea hydrophobic contribution, while the remaining portion are not. This may lead to a situation where the fluorescence will get its peak position at intermediate energies. But we do not believe that the mismatched duplex may exhibit a partial effect from the hydrophobic contribution. Fig. 4b summarizes these observations from the point of view of the nucleotides. The mismatched duplex, due to its sensitivity to environmental conditions via the unpaired hydrogen bond, responds to the presence of urea, a chaotropic agent that disrupts water structure, by occupying lower energy levels.

**Fig 4.**
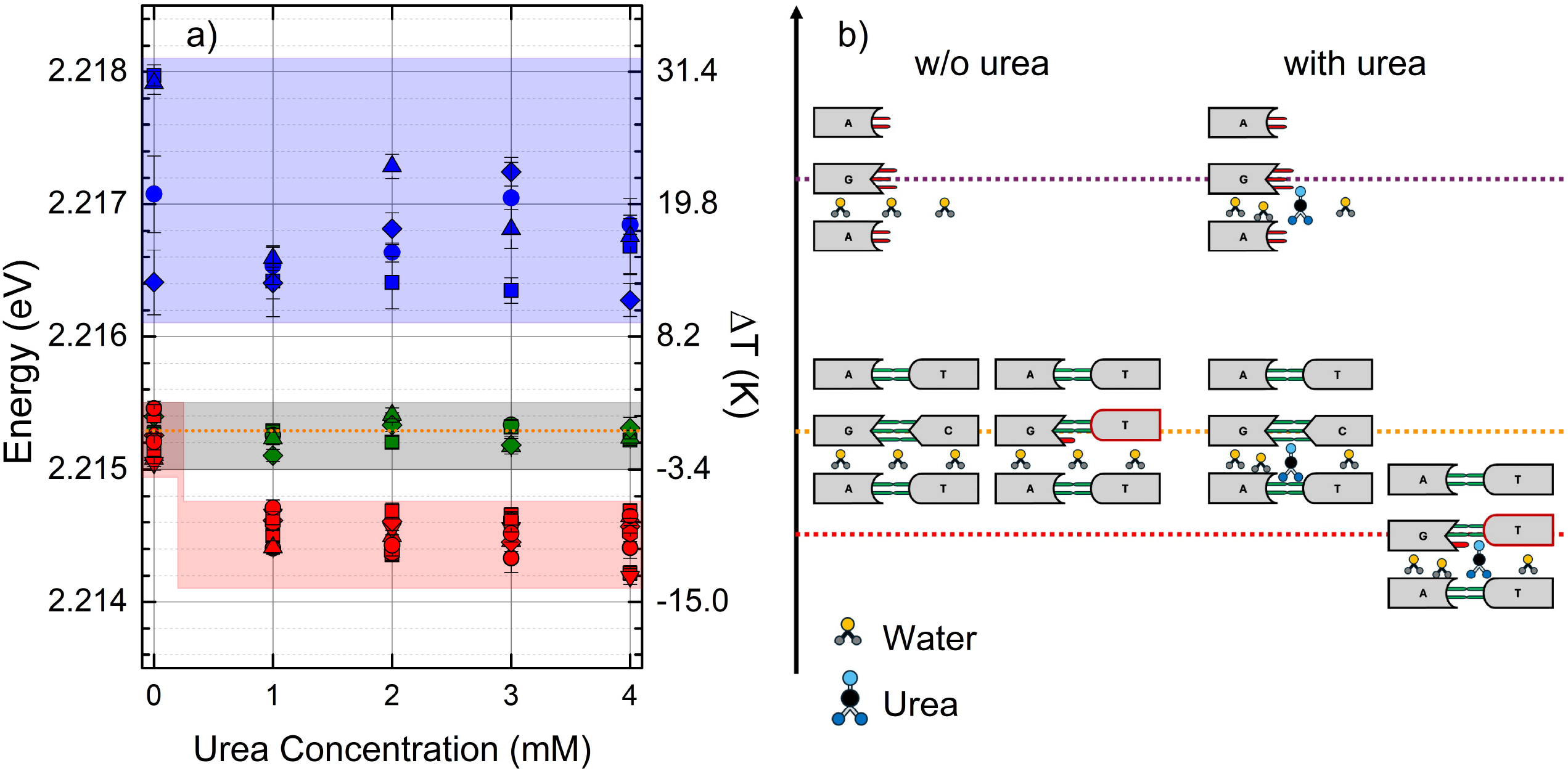
Energy diagram of the urea effect on the LMP in single- and double-stranded states, including mismatches. (A)Emission energy of the linear molecular probe (LMP) as a function of urea concentration, measured under three conditions: single stranded state with a scrambled target (blue area, blue data points), double stranded state with a perfectly matching target (gray area, green data points), and double stranded state with a target containing a single mismatched nucleotide (red area, red data points). Each data point is shown with its corresponding standard deviation. Shaded areas are provided to illustrate overall trends in energy change in increase of the urea concentration. However, we believe that for the red shaded area (double stranded state with mismatched nucleotide) the effect of energy reduction (red shift) has a binary character. **b)** energetic diagram of the LMP in single and double stranded states, where the double stranded state is formed with perfectly matching target and mismatched target (SNP is indicated by the nucleotide with a red rim). Disruption of the water structure by urea molecules results in lowering of the free energy of the mismatched duplex, making it energetically favorable even as the perfectly matched duplex – red shifted. Unpaired hydrogen bonds in the single-stranded and mismatched duplex states are shown in red.

## Conclusions

Early proposed method was validated using the LMP in both single and double stranded configurations. Measurement of the fluorescence emission energy of the LMP allows for a clear distinction between the single stranded state and the duplex states, where hybridization with either a perfectly matching or a single mismatched target induces a notable change in free energy. Under standard conditions, however, the method does not resolve the difference between perfectly matched and mismatched duplexes, as the energy difference between these two states is minimal. Involving a chaotropic agent into reaction, such as urea, perturbs the order of surrounding water molecules, thereby increasing the system’s entropy. This entropy increase leads to the decrease in free energy, enabling the separation of energy levels for the perfect and mismatched duplexes. The observed energy separation is attributed to the entropy of the single unpaired hydrogen bond present in the mismatched duplex.

In addition, the advantage of determining the LMP energy from fluorescence spectra, in difference to the methods using fluorescence intensity only, was clearly demonstrated. This approach overcomes limitations commonly associated with traditional techniques such as FISH and enables differentiation between duplexes containing polymorph nucleotides. Furthermore, it may offer insights into the molecular nature and origin of the molecular structures having such polymorph nucleotides.

## Author Contributions

Conceptualization: Oleh Fedorych.

Investigation: Marcello Gottschalk, Robin Jacobi, Oleh Fedorych.

Methodology: Sophia Rosencrantz, Oleh Fedorych.

Visualization: Oleh Fedorych.

Writing – original draft: Oleh Fedorych.

Writing – review & editing: Ruben R. Rosencrantz, Oleh Fedorych.

## Notes

### Competing Interest Statement

Oleh Fedorych is founder, shareholder and holds patent covering the method. Ruben R. Marcello Gottschalk, Robin Jacobi and Sophia Rosencrantz, have no competing interests to declare.

### Funding

- This work is co-funded by the Europäischer Fonds für regionale Entwicklung (EFRE).

