## Supporting information for "Formation of DNA duplexes in the presence of urea as a chaotropic agent"

**S1 Fig.** Fitted peak amplitudes of the linear molecular probe, averaged over 60 spectra, measured under three conditions: **a)** single-stranded state with a scrambled target (open blue squares), **b)** double-stranded state with a perfectly matching target (open black squares), and **c)** double-stranded state with a target containing a single mismatched nucleotide (open red squares). Each data point is normalized to a spectrometer integration time of 1 second and is shown with its corresponding standard deviation bars.
