## Supplementary figures and images for "Formation of DNA duplexes in the presence of urea as a chaotropic agent"

### S1_Fig1

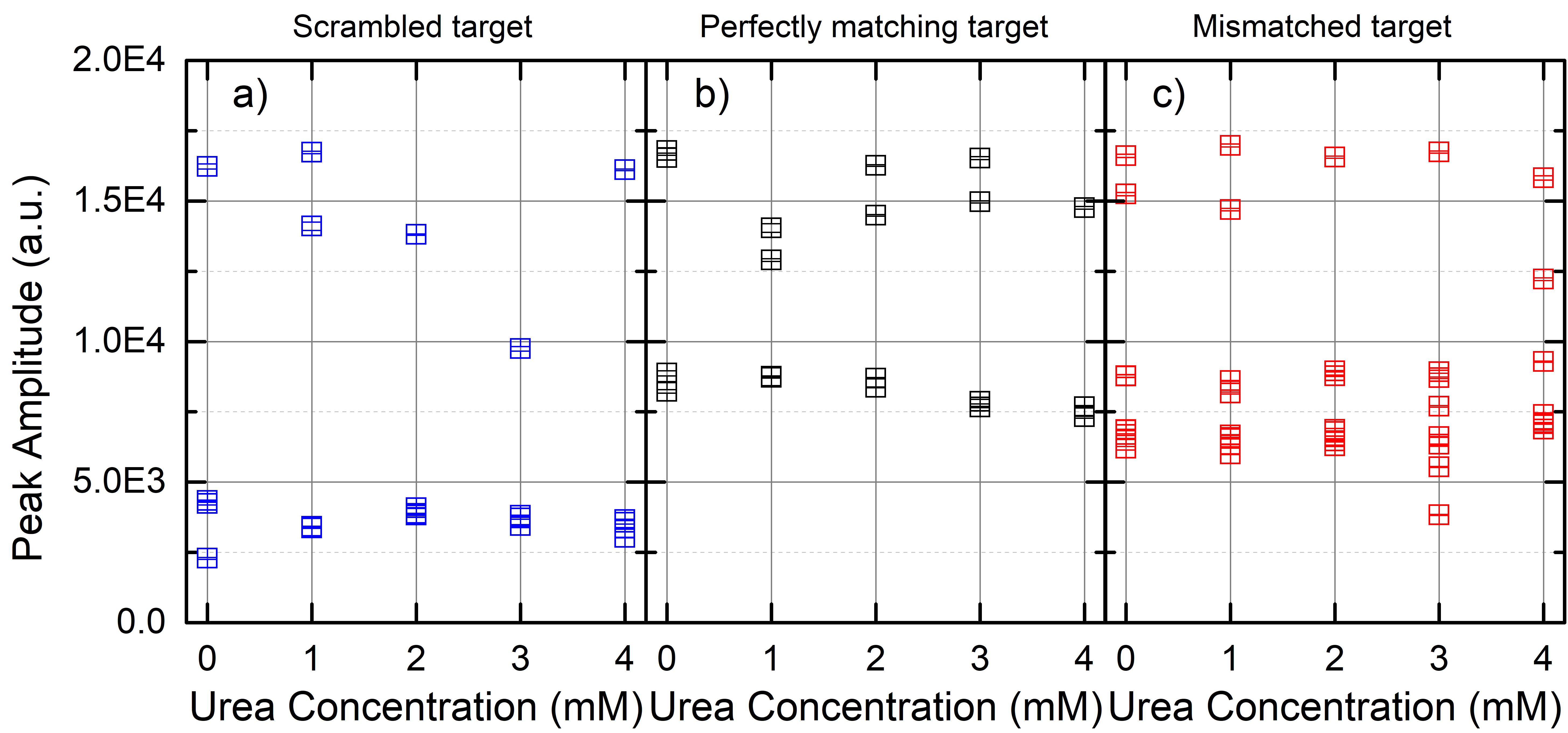
